# Parameter Optimization for Iterative MINFLUX Microscopy enabled Single Particle Tracking

**DOI:** 10.1101/2025.06.10.658837

**Authors:** Bela T.L. Vogler, Giovanni De Angelis, Ziliang Zhao, Christian Eggeling, Francesco Reina

## Abstract

MINFLUX fluorescence microscopy is a recently introduced super-resolution approach for studying cellular structures and their dynamics with highest detail. Iterative MINFLUX (*iMFX*) performs Single Particle Tracking (SPT) at *runtime*. The *ad hoc* signal interpretation necessary to sustain the method relies on several parameters, which need to be optimized in relation to the sample under study, such as fluorescent lipid analogues in membranes, to ensure the fidelity of the measurement. We propose a parameter optimization strategy, an overview of the most important parameters, present a theoretical upper limit for trackable diffusion rates, and demonstrate *iMFX*-enabled SPT of fast (⟨*D*_MSD_⟩ = 2.5*μm*^2^/*s*) lateral Brownian motion of lipids in membranes.

## Main

Microscopy-based Single Particle Tracking (SPT) is a popular method to investigate the molecular dynamics and interactions in living systems [1–3]. SPT is supported by a large variety of microscopy techniques [2, 4–6], including recently developed MINFLUX super-resolution microscopy [7, 8]. MINFLUX microscopy localizes single isolated fluorescent emitters by displacing an excitation beam with a central intensity minimum (i.e., a doughnut) in a pre-defined pattern, called Target Coordinate Pattern (TCP), around the initial position estimate of the emitter (see Methods). Imagining the discrete positions where the excitation beam is displaced as laying on a circular path, an important parameter for further calculation is the diameter of this circle, *L* (Figure 1ab) [7]. The commercially-available implementation of MINFLUX microscopy, hereinafter referred to as iterative MINFLUX (*iMFX*), generates the initial position estimate by searching for sources of fluorescent signal across the region of interest (Figure 1b) and then operates successive pattern scans at increasingly reduced *L* to refine the localization estimate (Figure 1cd) [8]. The lower limit of the localization precision derived by one TCP iteration, in the absence of noise, is determined by the Cramer-Rao Lower Bound (CRLB) of the maximum likelihood estimator used to locate an emitter within the TCP (see Supplementary Material of [7], p.8):

**Figure 1.**
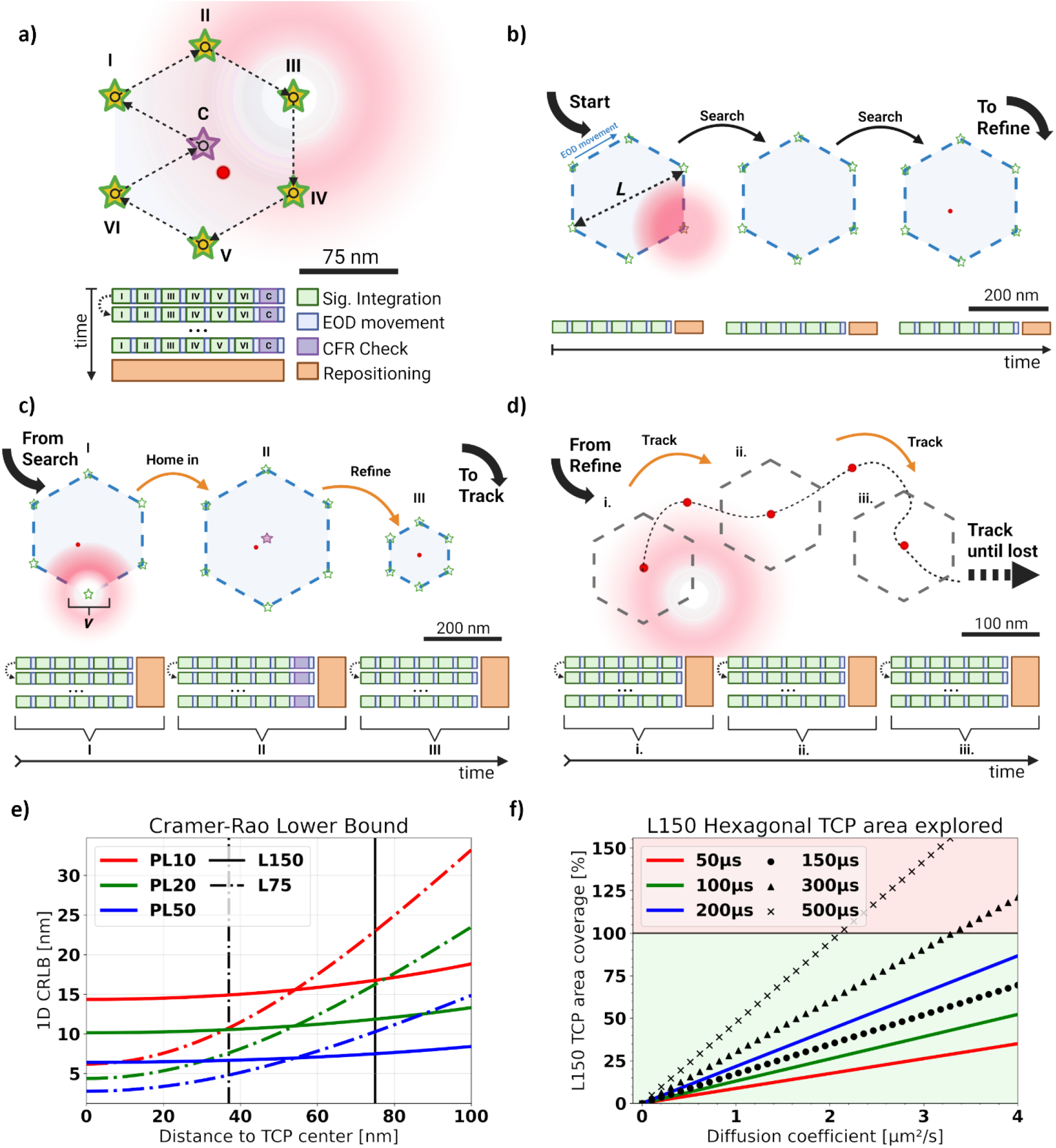
Principle and Theoretical Considerations of iMFX-enabled Single Particle Tracking. a) An illustration of a single *iMFX* TCP scanning pattern. The beam is steered along the marked path of vertices highlighted by green stars. The optional central scanning spot of the Center-Frequency-Ratio-Check (CFR-Check) is highlighted by a purple star. The numbers correspond to the sequence in which the scanning positions are reached. The EOD movement is represented by black arrows. The blocks on the bottom illustrate how much time it takes to execute each step of the TCP iteration. We represent the possibility of needing multiple repetition of the same iteration to obtain a localization by repeating the block diagram along the vertical axis. In all following panels the *Target Coordinate Pattern* (TCP) iterations follow this same scanning scheme. (*Created in BioRender. Vogler, B. (2025) https://BioRender.com/f7m5i5s*) b) Illustration of the grid-search process that finds candidate emitters in the sample. We highlight how this step is performed using no beam shaping but scanning a gaussian beam focus in a hexagonal pattern along pre-defined positions in the selected Region of Interest (ROI). (*Created in BioRender. Vogler, B. (2025) https://BioRender.com/vu4km6i*) c) After a candidate emitter is detected in the search stage, the localization algorithm refines the localization estimate through successive steps in which the diameter of the TCP, *L*, is progressively reduced. The capital roman numbers I-III correspond to the iterations as shown in Table 1. (*Created in BioRender. Vogler, B. (2025) https://BioRender.com/0ngjdaj*) d) Exemplification of the *iMFX*-enabled Single Particle Tracking process in which the last iteration of the sequence (III in panel c) is repeated and the particle followed until lost. The lower-case roman numerals are an aid to highlight the order in which these steps are executed. In a real *iMFX* experiment, the sequence will be repeated more frequently and as such the final iteration will not be repositioned by such large intervals, and this representation serves only as a visual aid. (*Created in BioRender. Vogler, B. (2025) https://BioRender.com/yn314ae*) e) Theoretical CRLB (Equation 1) for MINFLUX measurements executed with a doughnut-shaped beam against the distance of the localized particle to the center of the TCP for different photon limits (*PL*) and TCP diameters *L*. The vertical lines mark the TCP radius *L*/2 (solid line for *L* = 150*nm*, dashed one for *L* = 75*nm*). f) The fraction of a *L* = 150*nm* TCP area explored by a Brownian particle with constant diffusion rate *D* within different times-to-localization, i.e. the dwell time in addition to the hardware introduced temporal overhead. The colored solid lines correspond to the dwell-times used in this work. The background color corresponds to the respective area inside (green) and outside (red) of the TCP. This graph considers the time-to-localization, assuming that localizations are successfully estimated within a single TCP scan (⟨*η*⟩ = 1).

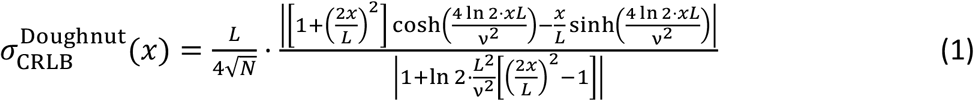

**Table 1.** Experimental MINFLUX Sequence Parameters for 2D Single Particle Tracking of Fluorescent Dye or QDot-labeled Lipid Analogues in the SLB –. (*) marks the parameters that were adjusted during the experiments.

| 2D Tracking | Pinhole Orbit [I] | 1<sup>st</sup> MFX Iteration [II] | 2<sup>nd</sup> MFX Iteration [III] |
| --- | --- | --- | --- |
| L (nm) | 284 | 302 | 150 |
| Pattern Shape | Hexagon | Hexagon | Hexagon |
| Photon Limit (counts) | 40 | 20 | 50 (*) |
| Laser Power Factor (times) | 1.0 | 1.5 | 2.0 (*) |
| Pattern dwell time (μs) | 500 | 100 | 200 (*) |
| Pattern repeat (times) | 1 | 1 | 1 |
| CFR Threshold | -1.0 | 0.5 | -1.0 |
| Background Threshold (kHz) | 15 | 30 | 30 (*) |
| MaxOffTime (control param.) | 3ms | 3ms | 3ms |
| Damping (control param.) | 0 | 0 | 0 |
| Headstart (control param.) | -1 | -1 | -1 |
| Stickiness (control param.) | 4 | 4 | 4 |

Here, *L* is the diameter of the TCP, *v* is the diameter (or full-width-at-half-maximum, FWHM) of the excitation beam, *N* is the number of photons collected, and *x* is the distance of the emitter to the center of the TCP.

For creating a single particle trajectory, *iMFX* necessarily performs particle position estimation and trajectory linkage at runtime. This is in stark contrast to conventional SPT applications, where analysis of this kind usually happens in post processing. Thus, whatever data is returned by the microscope is an interpretation of the signal produced by the sample based on the given acquisition parameters like *L, N* or *v*. This allows to seamlessly follow single emitters in time over all three dimensions of space with high spatiotemporal resolution [7, 8]. However, merging the *ad-* and *post hoc* stages leads to a significant increase in process complexity and *a priori* considerations, i.e. parameter choice. Counter to conventional SPT, maladjusted acquisition parameters drastically limit the ability to track emitters and can lead to premature interruption of tracking of a viable, i.e. not photobleached, target.

Here, we focus on engineering only the last iteration of the full sequence needed to obtain a valid localization as it is the most relevant for tracking applications. As per usual in SPT, the goal is to keep the time between consecutive localizations to a minimum as the particle is constantly moving, while maintaining a sufficiently low level of spatial uncertainty. To generate a localization, a set number of photons called the Photon Limit (*PL)* needs to be acquired within an array of consecutive TCP cycles *η*, which takes a total finite time-to-localization *t*_loc_, determined by *η*, the dwell time (*dT, t*_*dwell*_) and the hardware overhead time 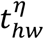, i.e. the delay introduced by the hardware response and execution time, which scales with *η* and the TCP geometry (see Methods). Therefore, the capability of *iMFX* to follow single emitters is significantly dependent on predefined parameters like *PL, dT* and the excitation laser power (which determines the frequency of photons *N* collected, i.e. how fast *PL* is reached), the time overhead 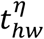 and sample-dependent variables such as the signal-to-noise ratio and the photophysical characteristics of the target fluorescent dye.

It is important to point out that during a localization estimation with *η* cycles, photons are only integrated for the time window 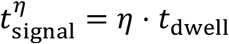, while the fluorescent target is moving for the entirety of *t*_*loc*_ (compare Equation 7). Therefore, both timeframes need to be considered separately. *t*_dwell_ experiences a lower limit based on the hardware response time that, given the *kHz* sampling frequencies of *iMFX*, cannot be assumed instantaneous

Assuming a Brownian particle with constant non-zero diffusion coefficient *D*, we can model the average distance traversed by the particle during a localization estimation as the standard deviation 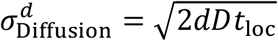 of the probability distribution of particle localizations [9], where *d* is the number of spatial dimensions and *t*_loc_ is the time-to-localization elapsed. While the position of the TCP center is updated after each successful localization, it remains unchanged during the cycles, i.e. during the estimation. Assuming one-dimensional diffusion, we can therefore estimate the average particle distance *x*, from Equation 1 to the TCP center by the 1D standard deviation 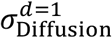, i.e. 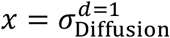. Therefore, the CRLB (Equation 1) depends on *t*_loc_ and on the particle diffusion rate *D*, and thus the CRLB is categorically larger for tracking experiments compared to imaging i.e. localization of fixed emitters (Figure 1e). This effect is more dramatic when, for instance, the diameter of the TCP is reduced in an attempt to increase the precision of localization without considering other parameters. Therefore, a delicate balancing act is necessary to bring together a reliable tracking experiment using *iMFX*, where expectations of the diffusion rate, as well as the photophysics of the fluorescent label and relevant scanning parameters, namely *PL, dT* and *L*, must all be taken into consideration. This is not as problematic in the case of immobile particles (*D* = 0), since in these cases it is possible for *iMFX* to ideally perfectly center the TCP over the emitter and optimization is required only to match the scanning parameter and the photophysics of the fluorescent label. This corresponds, in Figure 1e, to the values of the CRLB where the distance from the TCP center is zero.

The *iMFX* position estimation algorithm consists of two stages: 1) the initial position estimation for a detected emitter using a least mean squares estimator, and 2) the unbiasing step that corrects the initial estimate and relies on a set of pre-determined numerical coefficients [8]. The coefficients necessary to the unbiasing step are provided with the instrument and are the numerical solution to an optimization problem involving *PL* and *L*. Therefore, the *PL* must be carefully adjusted when attempting to increase the photon output of the target, for example by employing brighter fluorescence emitters, longer dwell times or higher excitation laser power [10]. In order to guarantee a reliable position estimate, the *PL* parameter in the MINFLUX sequence file should be matched with the expected average photon count (⟨*N*⟩) per unit time 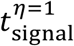. It is, in fact, desirable to set the *PL* to a slightly underestimated value for the emission of the target fluorophore. This will ensure that each TCP iteration will lead to a successful localization, therefore maximizing the sampling efficiency in terms of minimizing the time-to-localization. However, if the photon limit is set too low, the localization algorithm does not take full advantage of the photon budget afforded by the target fluorophore. This is because the *iMFX* localization estimator is fixed *a priori* compared to the measurements and is optimized to produce localization with *PL* photons. While a slightly higher number of actually detected photons compared to *PL* can still prove beneficial, significantly larger deviations will cause errors during the unbiasing step [11]. These errors get more severe for larger *L*, as the approximation of the central minimum of the excitation beam (i.e., the doughnut) by an ideal parabola (as done in practice) does not hold in that case, thereby causing a larger bias in the estimator.

Given that emitters are only tracked while inside the TCP, there is a risk for them to escape should the area 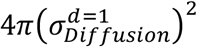, which is explored by an emitter during the time-to-localization *t*_loc_, approach the TCP area π*L*^2^. This solidifies an upper limit for trackable diffusion rates *D* given a specific *t*_loc_ (Figure 1f). If a particle expresses a diffusion rate beyond this limit, *iMFX* will not be able to produce reliable localizations, if any at all. From Equation 1 it becomes clear that while the spatial resolution is increased linearly by reducing *L*, the risk of losing the emitter rises quadratically.

As a reminder, during position estimation, the excitation beam is successively centered on the scouting spots, i.e. the TCP vertices. Here it remains for a fraction of the *dT* during which the fluorescence signal is integrated. While this strategy is ideal for an immobile target, it poses a fundamental issue of SPT experiments. Since an *iMFX* assesses the position of the tracked particle at the end of the TCP cycle, given sufficient photons have been collected, it cannot refine the estimate based on the evolution of the signal during the observation. This is, however, not necessarily desirable, since such a procedure would further increase the overhead time between observations.

From our theoretical considerations we know that the excitation laser-power, *PL*, and *dT* are the most impactful parameters to consider when optimizing the *iMFX* time-to-localization (see Methods). While the choice of laser-power strongly depends on the sample used (e.g. the brightness or photobleaching properties of the target fluorophore) both *PL* and *dT* can be chosen arbitrarily before the experiment. Consequently, we experimentally investigated the influence of these two parameters on *iMFX*-enabled SPT measurements. To this end, we performed SPT of biotinylated lipid analogues (DSPE-PEG-Biotin, tagged with streptavidin-coated Quantum Dots (QDs)) embedded in a homogeneous fluid continuous Supported Lipid Bilayer (SLB, 100 % unsaturated 1,2-Dioleoyl-sn-glycero-3-PC (DOPC) lipid) [12] (Figure 2a). We intentionally employed large (≈ 15*nm* − 20*nm* in diameter [13]) and bright metallic core QDs as luminescent tags to slow down the diffusion of the target biotinylated lipid analogues while preserving their characteristic free diffusion [14–17]. The employed SLBs were either generated by lipid depositions in enclosed flow-chamber systems or are Giant Unilamellar Vesicle (GUV) patches. We performed the main part of our experiments on GUV patches, as they offer an easy and well-explored model system.

**Figure 2.**
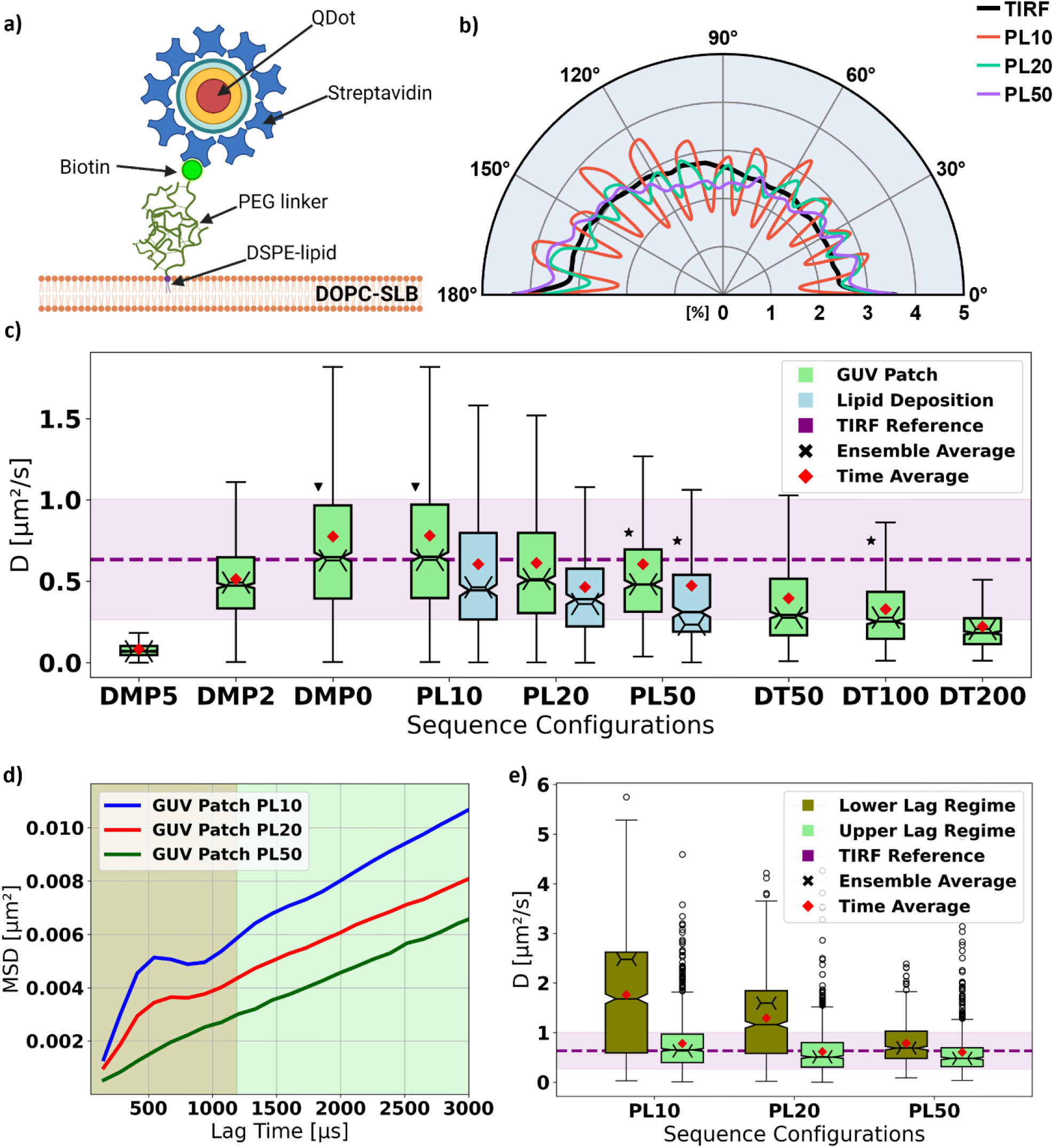
Experimental iMFX-enabled Single Particle Tracking (SPT) of QD-labeled lipid on a Supported Lipid Bilayer (SLB) Dependencies on hardware parameters. a) Biotinylated lipid analogues (DSPE-PEG-biotin) tagged with streptavidin-coated metallic core quantum dots (QDs) were incorporated in homogeneous DOPC SLBs and diffused freely in the membrane. b) Spline interpolated distribution of turning angles between successive displacement vectors of different ensembles of lipid trajectories, detected using *iMFX*-SPT with different photon limits (PL) and the same TCP diameter (L=150nm). As a reference, we reported the same distribution obtained using SPT experiments performed on a custom TIRF setup. c) Comparison of the measured diffusion coefficients across different sequence configurations tested on SLBs generated using GUV patches (green) or flow chamber lipid deposition (blue): different damping parameter *DMP* (with *PL* = 10 and *dT* = 100), photon limit *PL* (with *dT* = 100 and *DMP* = 0), and dwell time dT (with *PL* = 50 and *DMP* = 0) settings. Each boxplot (center line, median; red diamond, mean; box limits, first to third quartile; whiskers, 1.5x interquartile range; notch, height proportionally to interquartile range) corresponds to the distribution of diffusion coefficients extracted from the detected trajectories using the OLSF routine on the *Upper* Lag Regime of the MSD curves. We highlight the time-average (red diamond) and ensemble-average (black cross) diffusion coefficients for each dataset. As reference we provide the distribution of diffusion coefficients (average diffusion coefficient (purple line) and standard deviation (lilac-shaded area) determined) of the TIRF experimental data on a QD labelled GUV-patch SLB using the same analysis routine. Given the equality of time-average and ensemble-average diffusion coefficient for this dataset, we chose to display only one line. Each triplet of datasets focusing on a specific parameter, e.g. *PL*, has been taken on separate samples with the exception of the GUV Patch *DMP* and *PL* sets, which share the sample. Two datasets have been taken on the same sample with the same sequence (*PL* = 10, *DMP* = 0, *dT* = 100*μs*; black triangle), while three other sets have been taken on different samples with the same parameters (*PL* = 50, *DMP* = 0, *dT* = 100*μs*; black star). d) Illustration of how different values of *PL* contribute to generating the two Lag Regimes observed in our SPT experiments using the ensemble average MSD curves calculated from our GUV-patch SLB experiments. The shading highlights the two regimes (Lower Lag Regime: olive, Upper Lag Regime: light-green). We use an arbitrary threshold of 9 lags (≈ 1190*μs*) to separate between the regimes. e) Comparison boxplot (center line, median; red diamond, mean; box limits, first to third quartile; whiskers, 1.5x interquartile range; notch, height proportionally to interquartile range; points, fliers) of the extracted diffusion coefficients (as in c) for the GUV-patch SLB experiments between the Lower and Upper Lag Regimes, together with the reference values from the TIRF experiments (as in c).

Table 1 lists all relevant experimental *iMFX* sequence parameters. With *L* = 150, Stickiness = 4 and MaxOffTime = 3*ms* (see Appendix 2, Parameter Overview), we in our experiments chose to stay lenient during tracking to catch as many datapoints as possible, which gave us more freedom in data analysis but required *post hoc* artifact removal.

In our analysis, we split trajectories whenever the number of cycles *η* between consecutive localizations exceeded *δ* = ⟨η⟩ + std(η), which effectively removed large time-gaps, mid-trace-particle-swap events, and jitter. Further, using our previous SPT analysis pipeline, we eliminated artifacts due to e.g. label-induced cross linking of various lipids [18]. We then finally extracted the lateral diffusion coefficient *D*_MSD_ and the dynamic localization uncertainty σ_MSD_ from a custom Mean Squared Displacement (MSD) implementation suited for inhomogeneous time-lags using Optimal Least Squares Fitting (OLSF) [19]. As a reference for the obtained values of *D*_MSD_, we performed SPT on the same QD-labelled lipids in a similar SLB using a custom-built microscope with Total-Internal-Reflection Fluorescence (TIRF) excitation and camera-based detection [20], and analyzed the data using the same pipeline.

We first tested possible bias in the *iMFX* data by checking whether the tracks showed an isotropic progression as expected for Brownian diffusion. The Turning Angle Distribution (TAD) demonstrates an additional geometric bias in that a set of directions is preferred with decreasing *PL* (Figure 2b, *dT* = 100*μs*) digitizing the direction of positional updates, i.e. too low values of *PL* resulted in a biased detection of diffusion. This is a direct result of the reduced number of acquired photons and the thus reduced size of the statistical data sample available for analysis.

Before continuing the investigation of the influence of *PL* and *dT*, we inspected another experimental *iMFX* parameter, the galvo damping *DMP* (see Appendix 2, Parameter Overview). Therefore, we reviewed values of *D*_MSD_ obtained for different *DMP* (Figure 2c left side, *PL* set to 10). Since *DMP >* 0 led to an underestimation of particle motility, we generally kept the galvo damping off (*DMP* = 0) in our experiments.

As mentioned above, our isotropic progression analysis highlighted a favorable use of larger *PL* values. Unfortunately, we had to reduce *PL* to avoid missing faster diffusion events, i.e. to sample a higher bandwidth of particle diffusion rates, highlighted by an artefactual dependency of the extracted values of *D*_MSD_ on *PL* (Figure 2c): While higher *PL* entail a smaller localization uncertainty σ_MSD_, i.e. an in principle higher spatial precision (Table 2), the determination of values of *D*_MSD_ became biased towards slower moving emitters, i.e. emitters moving too fast were lost and the reported values of *D*_MSD_ appeared to underestimate the reference value of the TIRF-SPT experiments (Figure 2c, Table 2). We observed the same characteristics for both kinds of SLB-sample preparation. The coinciding change of σ_*MSD*_, *D*_MSD_ and *PL* can be traced back to the CRLB’s dependence on the number of photons acquired and the aforementioned increased possibility of losing particles moving too fast for too long times (*t*_loc_), i.e. the time required in total for a localization (Figure 1e,f).

**Table 2.** Experimental Results Reveal Optimization Tradeoff Between Trackable Diffusion Rates and Localization Error for iMFX-enabled Single Particle Tracking. *–* In addition to the ideal CRLB 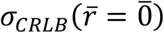 (see Supplementary Material of [7], p.11) we show the mean of the apparent lateral diffusion coefficient *D*_*eMSD*_ as well as the lateral localization error σ_*eMSD*_ extracted using Optimal Least Squares Fitting (OLSF) on the ensemble average Mean-Squared-Displacement (*e*MSD) assuming momentary Brownian motion. Here *η* refers to the number of cycles, *t*_*loc*_ is the time-to-localization, *N* is the number of photons, *N*_*PL*_ is the *PL* and ⟨*…* ⟩ the scale-appropriate average value. Shading indicates sample reference Dye on GUV-patch SLB (orange), QDs on GUV-patch SLB (green), QDs on Lipid-Deposition-SLB (blue) and the TIRF reference (lilac).

| PL | DMP | DT | $D_{eMSD}$<br>[$\mu m^2/s$] | $\sigma_{eMSD}$<br>[nm] | $\sigma_{CRLB}$<br>[nm] | $\langle \eta \rangle$ | $\langle t_{loc} \rangle$<br>[$\mu s$] | $\frac{\langle N \rangle}{\langle \eta \rangle N_{PL}}$ |
| --- | --- | --- | --- | --- | --- | --- | --- | --- |
| 10 | 0 | 100 | 2.46 | 30.30 | 20.03 | 1 | 150 | 2.13 |
| 10 | 5 | 100 | 0.07 | 16.52 | 20.03 | 1 | 150 | 4.10 |
| 10 | 2 | 100 | 0.49 | 15.53 | 20.03 | 1 | 150 | 4.49 |
| 10 | 0 | 100 | 0.63 | 28.02 | 20.03 | 1 | 150 | 3.36 |
| 10 | 0 | 100 | 0.63 | 27.96 | 20.03 | 1 | 150 | 3.36 |
| 20 | 0 | 100 | 0.52 | 22.78 | 14.35 | 1 | 151 | 2.27 |
| 50 | 0 | 100 | 0.48 | 14.05 | 9.08 | 2 | 282 | 0.71 |
| 10 | 0 | 100 | 0.46 | 30.37 | 20.03 | 1 | 150 | 1.53 |
| 20 | 0 | 100 | 0.36 | 23.51 | 14.35 | 2 | 281 | 0.67 |
| 50 | 0 | 100 | 0.23 | 13.46 | 9.08 | 5 | 676 | 0.22 |
| 50 | 0 | 50 | 0.28 | 4.67 | 9.08 | 5 | 437 | 0.23 |
| 50 | 0 | 100 | 0.28 | 5.50 | 9.08 | 3 | 414 | 0.42 |
| 50 | 0 | 200 | 0.21 | 10.91 | 9.08 | 2 | 483 | 0.82 |
| TIRF SPT |  |  | 0.64 | 39.86 | — | 1 | 14925 | — |

We next investigated the effect of different *dT* on the experimentally detected diffusion rates, while maintaining *PL* = 50 (Figure 2c). Confirming Equation 5, we revealed larger average values of *D*_MSD_ for smaller *dT*, reaching the reference value of the TIRF-SPT experiments only for very small *dT* ≤ 50*μs*. Yet, one has to keep in mind that that due to the additional hardware-induced overhead time 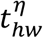, there was a lower limit to reducing *dT*. Given that photon emission follows Poisson statistics and assuming a fixed average emission rate ⟨θ_p_⟩, the average number of detected photons per cycle 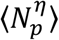 is linearly dependent on the time window of signal integration (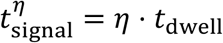, see above). As a consequence, for lower *dT* a larger average number of TCP cycles ⟨*η*⟩ is required to collect a sufficient number of photons (*PL*) per successful localization (compare Table 2). This again results in a larger or equal time-to-localization *t*_loc_ (compare Equation 4) given that each TCP cycle causes additional hardware overhead. As a reminder, *t*_loc_ is determined by the number of TCP cycles *η* times the dwell time *dT* plus the overhead time 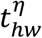 (Equation 7).

In a nutshell, for a given ⟨θ_*P*_⟩, fewer and shorter cycles per localization mean a higher frequency of position updates, which permits following faster particles at a reduced distance from the TCP center, thereby minimizing the localization error induced by the particle movement, i.e. the offset between the actual (and unknown) particle position and the estimation produced. This strategy effectively improves the overall spatio-temporal resolution. Improving the localization precision by collecting more photons over a longer integration time (*t*_signal_) may not always be viable, since increasing *t*_*loc*_ in turn increases the deviation between estimation and actual particle position or renders the position estimation impossible altogether. This is due to an increase in the area explored by the diffusing particle during the photon collection (compare Figure 1f). Additionally, it may seem a plausible approach to increase the temporal resolution by adopting TCP scanning patterns with fewer positions. This has already been employed in the past [7, 8], and it does indeed marginally increase the sampling rate, since fewer movements of the laser must be performed during one TCP, and thus less time is spent on one acquisition cycle. As we have shown (Figure 2b), this again results in an angular bias in the position estimator, which may be undesirable. Therefore, adopting a hexagonal TCP is preferrable, at least when probing diffusive motion with a high degree of randomness[8].

While calculating the ensemble-MSD from our experiments, we noticed that they expressed two distinct discontinuous linear regimes, most prominently for low values of *PL*, i.e. high temporal resolutions (Figure 2d). Though varying in severity, we found the distinct *kink*-like shape in all of our fastest datasets. Consequently, for the entirety of this work, we distinguish between the *Lower* Regime MSD, meaning the part with or prior to the kink-like shape, and the *Upper* Regime, referring to the continuous linear part beyond that. Here, the words *Lower* and *Upper* references the interval of time lags *τ* in which discontinuity appears. We noticed a correlation between the severity of the kink and the employed *PL* (Figure 2d). Given that a lower *PL* enables faster sampling (see Methods and Table 2), we further noted a correlation between the sampling frequency of *iMFX* (∼1/⟨*η*⟩) and the severity of the discontinuity of the MSD curve. The same was true for the dynamic localization error σ_MSD_, which got worse (i.e. increased) with increased sampling frequency (Table 2).

We further anticipated explaining the two-regime characteristics of our experimental MSD curves. We safely excluded the possibility of our tracked particles experiencing strong confined or compartmentalized diffusion, since our membrane bilayers were very fluid, consisting of one type of unsaturated lipid only. Therefore and following previous work [21], it is most probable that the *Lower* Regime MSD did not result from the diffusive motion of the particle but was rather due to the wiggling motion of the QDs (≈ 15*nm* − 20*nm* in diameter [13]) attached to the PEG(2000) linkers (≈ 3*nm* in diameter [22]) on top of the particle (comparable to [21] Figure 5). This became apparent when comparing the extracted values of *D*_MSD_ for both *Lower* and *Upper* Regime (Figure 2e). On one hand, the *Upper* Regime exhibited diffusional rates close to those of our TIRF microscopy reference experiments, and appeared only marginally dependent on *PL*, highlighting the true mobility of the lipid. On the other hand, *D*_MSD_ values of the *Lower* Regime were above those of the references (Figure 2e). Therefore, we considered the *Upper* Regime as the region of interest for our work. Hence, throughout this work, except for Figure 2e, we list only the diffusion rates extracted from the *Upper* MSD Regime (Figure 2c, 3b, Table 2) disregarding the *Lower* Regime. Further investigations into the cause of the MSD discontinuity in general were beyond the scope of this work.

Given that Brownian motion is a homogeneous, isotropic and memoryless process, particle systems exhibiting Brownian motion are expected to be ergodic, i.e. to have an ergodic ratio *ε*_Brownian_ = 1. Therefore, it is important to point out that all MINFLUX datasets, which reported diffusion coefficients close to the values of the TIRF microscopy reference, were characterized by an ergodic ratio of about *ε*_*QD*_ ≈ 0.8 as compared to *ε*_TIRF_ = 1.0 for TIRF. Following the above discussion, the observed ergodicity break was most likely due to the superposition of the wiggling motion of the QDs and the particle diffusion in the SLB.

*iMFX* interprets fluorescence signal at *runtime* to track individual emitting particles effectively in a single convenient process. As pointed out, it is necessary to optimize experimental parameters with great care and tailor them to the sample, which enables faithful reporting of the population of diffusing particles by the *iMFX* measurements. As listed above, this can be achieved by adjusting the TCP geometry and diameter *L*, the excitation laser power (according to the brightness and photostability of the chosen fluorescent emitter) but most importantly through *PL* and *dT*. Summarizing our theoretical and experimental considerations, it is evident that minimizing the time between consecutive localizations, while keeping *PL* as high as possible, is key to enabling high-fidelity *iMFX-*enabled SPT measurements. Given the theoretical definition of *t*_*loc*_ (Equation 5), a continuous fluorescent signal and any given *PL* and *dT*, the time-to-localization minimizes exactly when:

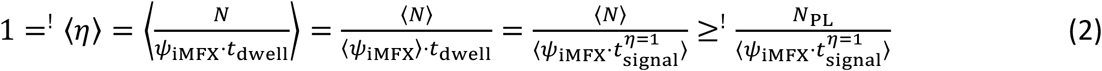

where ⟨*ψ*_iMFX_⟩ is the average photon detection frequency per localization (sometimes referred to as EFO), 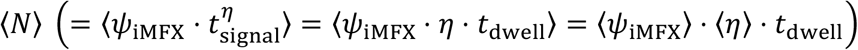 the average number of collected photons (sometimes referred to as ECO), *t*_dwell_ the dwell time per localization round as put in the sequence, and *N*_*PL*_ the photon limit.

For simplification, we may set the Brownian diffusing emitters to exhibit a constant average emission rate ⟨θ_p_⟩ during an optimized photon acquisition (⟨*η*⟩ = 1) and assume a similarly constant average photon detection rate ⟨*ψ*_iMFX_⟩ ≤ ⟨θ_p_⟩. In this case, we can give an estimation of the upper limit of possibly trackable 2D diffusion rates 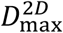 under the condition that the particle has to remain within the area of the TCP during the localization process (Methods).

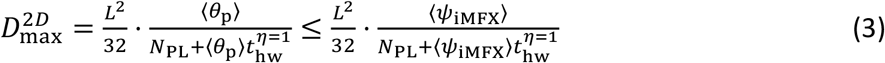

Prior to the estimation of a theoretical upper limit, we conducted *iMFX* enabled SPT experiments on fluorescent lipid analogues (Dipalmitoylphosphatidylcholine tagged with Abberior STAR-Red via a PEG-linker, DPPE-PEG) embedded within a GUV-patch derived SLB (70% DOPC and 30 % cholesterol (CHOL); Figure 3a). The addition of cholesterol was made to marginally slow down the diffusion (to ensure to stay within 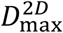) while maintaining its characteristic isotropic and unconfined movement. Following the same routine as before (with *PL* = 10, *dT* = 100*μs, DMP* = 0), we calculated the MSD. From its *Upper* Regime, we obtain an ensemble and time-average diffusion coefficient *D*_Dye_ ≈ 2.5*μm*^2^/*s* and an ergodicity coefficient of *ε*_Dye_ ≈ 0.98, i.e. close to perfect homogeneous diffusion (Figure 3b, Table 2).

**Figure 3.**
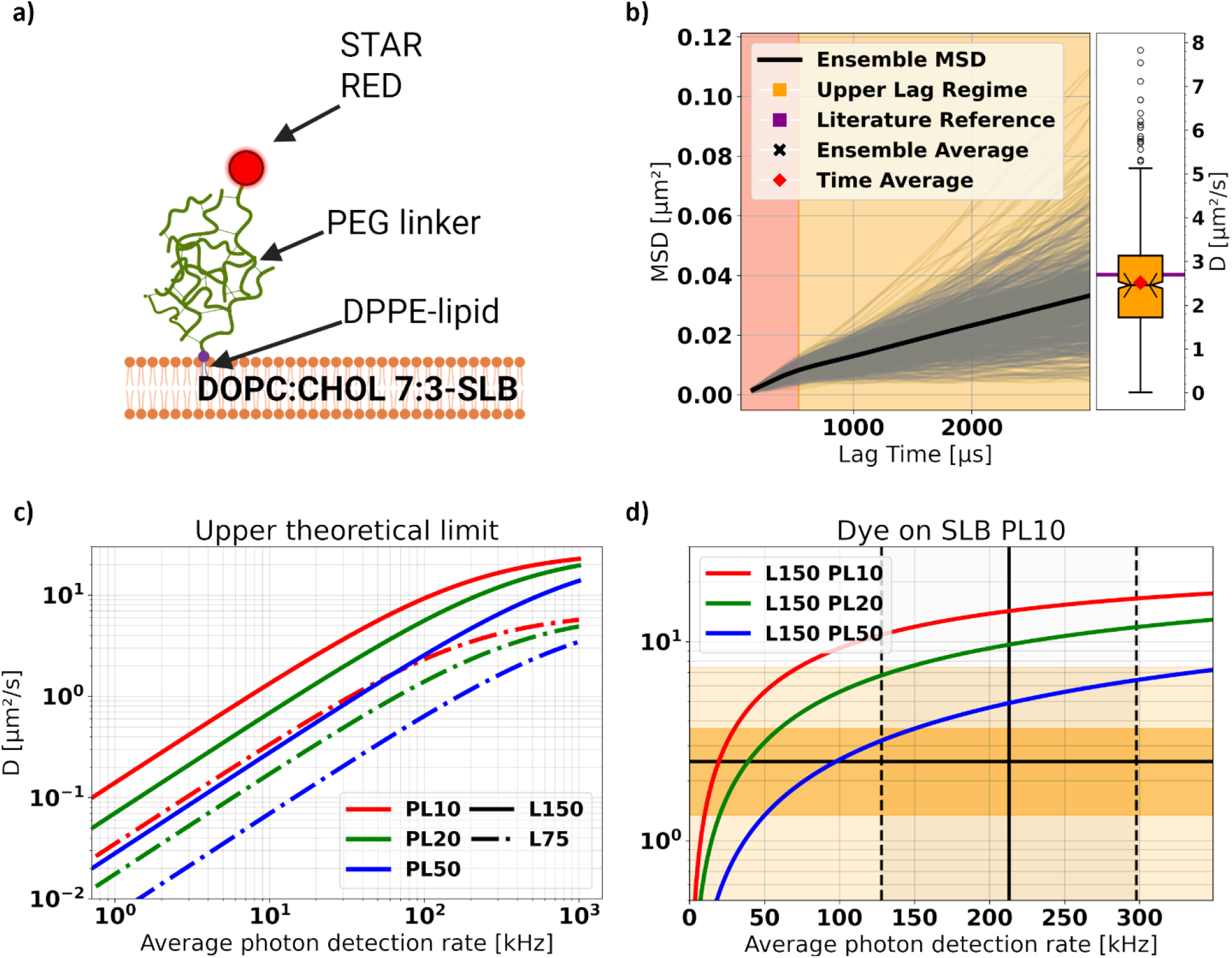
Theoretical and Experimental Considerations Reveal Upper Limit of Trackable Diffusion Rates. a) Fluorescent lipid analogues (DPPE-PEG2000-STAR RED) tagged with Abberior STAR RED were incorporated in homogeneous DOPC:CHOL 7:3 bilayers. b) Single-molecule MSD curves (gray lines), and the ensemble average MSD curve (black line) calculated for trajectories of the fluorescent lipid analogues diffusing in the SLB illustrated in a). The shading corresponds to the Lower (orange-red) and Upper (orange) Lag Regimes, with an arbitrary threshold set at 5 lags (≈ 670*μs*). The corresponding distribution of diffusion coefficients (time-average (red diamond) and ensemble-average (black cross)) extracted for the Upper Lag Regime of the MSD can be found as a boxplot (center line, median; red diamond, mean; box limits, first to third quartile; whiskers, 1.5x interquartile range; notch, height proportionally to interquartile range; points, fliers) to the right. The horizontal purple line marks a system specific reference-value taken from literature [23].c) Upper theoretical limit of trackable diffusion coefficients *D* against the average photon detection rate of the particle within one TCP iteration as calculated from Equation 3, in the absence of noise. Different line styles and colors represent unique TCP diameters *L* and photon limits *PL* as labeled. d) Zoom-in of the graph in c) for values of *D* and average photon detection rate more common in experimental practice. The black horizontal line indicates the ensemble (and time) average diffusion coefficient *D* for the dataset in panel b), while the vertical one represents the average frequency of photon detection ⟨*ψ*_*iMFX*_⟩. The shaded dark orange area represents the standard deviation of the distribution of *D* from panel b), whereas the lighter orange represents the full distribution. The vertical dashed lines represent the standard deviation of the photon detection rate.

In comparison to the QD-tagged samples, we noticed that the kink-like shape of the ensemble-MSD, though retained, was significantly more subtle. This implies reduced wobbling of the PEG-linker tagged with the dye rather than with the QDs, which was reflected in the contrast of ergodicity coefficients *ε*_*QD*_ ≈ 0.8 and *ε*_Dye_ = 1.0 for the same sequence parameters (i.e. *PL* = 10, *dT* = 100, *DMP* = 0). Consequently, *iMFX* successfully observed the expected isotropic Brownian Motion with a diffusion rate close to values reported in relevant literature [23].

We then compared our experimental results to theory. Figure 3c plots corresponding diffusion rate limits *D*^2*D*^ for different values of *L, PL* and ⟨*ψ* ⟩, highlighting the best chances to track fast moving emitters for large *L* (e.g. 150 nm) and ⟨*ψ*_iMFX_⟩ and low *PL* (e.g. 10). With photon emission frequencies in the range of 125-300 kHz (as in our experiments), *iMFX* should then be able to track emitters moving as fast as 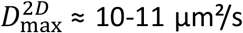 with optimized parameters (Figure 3d). Our experimental data of Figure 3b (*D*_Dye_ ≈ 2.5*μm*^2^/*s*, taken with PL10 and dT100) was thus well within these theoretical boundaries (Figure 3d). Given that in these experimental settings slightly more than *N* = 20 photons were detected on average per localization and cycle (*PL* = 10, first row and last column Table 2), resulting in an average photon detection frequency per localization ⟨*ψ*_iMFX_⟩ = ⟨*N*/*ηt*_*dwell*_⟩ of around 213 kHz (⟨*t*_loc_⟩ = 150 *μs*, Table 2 and Equation 3), one might argue that increasing the *PL* to 20 accordingly could prove beneficial in terms of localization precision while maintaining a rather faithful representation of the average diffusion rate. Further increase of the *PL* or *dT* or decreasing *L* for that matter would however result in an underrepresentation of the particle motility within the system. It is important to note that we always assume ⟨*η*⟩ = 1, which implies *t*_*dwell*_∼1/⟨*ψ*_iMFX_⟩ and *N*_PL_ ∼⟨*ψ*_iMFX_⟩ (compare Equation 3).

By theoretical and experimental means, we have demonstrated the implications of the iterative localization procedure of *iMFX*-enabled SPT and highlighted the resulting critical dependence on pre-defined acquisition parameters such as *L, PL*, and *dT* and deducted an upper limit of reliably trackable diffusion rates. We introduced a parameter optimization scheme (Appendix 1, *Iterative-MINFLUX Sequence Optimization Guide for 2D SPT Experiments*) to allow for the capture of a broader bandwidth of fast diffusion events and provided an overview of the most relevant *iMFX-* Sequence parameters (Appendix 2, *Overview of the Most Influential Sequence Parameters in Iterative-MINFLUX-enabled Single Particle Tracking*) alongside practical tips and warnings for *iMFX*-sequence manipulation. Finally, we experimentally demonstrated the possibility of reliable *iMFX*-enabled SPT of fast (⟨*D*_MSD_⟩ = 2.5*μm*^2^/*s*) lateral Brownian diffusion of fluorescent lipid analogues in model membranes.

Even though we have restricted ourselves to two-dimensional experiments for the sake of simplicity, our discussion and findings can be applied to the three-dimensional case given they share the same *modus operandi*. We should point out that all the considerations for Brownian diffusing particles in this text apply also to cases of directed or processive motion, such as those considered in [24, 25], since the instrument adopts the same detection strategy irrespective of the sample or the mode of motion thereof.

## Methods

### Continuous Supported Lipid Bilayer Composition and Formation

The continuous supported lipid bilayer (SLB) used as a target sample was composed of DOPC (1,2-dioleoyl-sn-glycero-3-phosphocholine, *Avanti Polar Lipids, Inc*.) with the addition of 0.01 Mol% of the fluorescent lipid analogue DOPE-ATTO488 (*ATTO-TEC GmbH*) for ease of identification under confocal fluorescence imaging. The targets of SPT were DSPE (1,2-distearoyl-sn-glycero-3-phosphoethanolamine) lipid analogues with a PEG(2000)-Biotin linker (*Avanti Polar Lipids, Inc*.), added in 0.15 ⋅ 10^−3^ Mol% to the lipid solution above. The SPT targets were labelled with streptavidin-coated metallic core QDs, whose fluorescence emission spectrum manifests as a symmetric peak centered around 655 nm (QDs 605, *ThermoFisher, Inc*.).

The continuous SLB production was based on a previously reported solvent-assisted lipid bilayer (SALB) formation method [12]. Lipids were first dissolved in chloroform upon arrival and kept as stock solutions at -20 °C. Before use, the desired amount of lipid stock solution was blow dried by a gentle stream of nitrogen gas and the lipid film was resuspended in isopropanol (IPA) at a concentration of 0.5 *mg*/*ml*. Glass coverslips (Epredia, 0.16 − 0.19*mm*, 26 × 76*mm*) were rinsed sequentially with ethanol, ultra-pure water and cleaned with detergent, they were treated with a plasma cleaner (Zepto One from *Diener Electronic GmbH, Plasma-Surface-Technology*) for 1 min before the assembly of the microfluidic flow channel with *Ibidi GmbH* sticky-Slide I Luer (0.1*mm* channel height), the flow channel was later connected to a *Hamilton, Inc*. gas tight glass syringe (2.5 *ml*) via appropriate connectors and tubing. A high precision syringe pump (*CETONI GmbH* Nemesys S) was used to control the liquid exchange. The flow channel was initially filled with PBS buffer, then the buffer was replaced with IPA at a flow rate of 50 *μl*/*min* for 5 *min*, target lipid/lipid mixtures in IPA was then introduced into the channel at 50 *μl*/*min* for 2 *min* and with incubation on the coverslip for another 5 − 10 *min*, finally PBS buffer was again introduced into the flow channel at 50 *μl*/*min* for 2 *min* for complete formation of continuous SLB, and with a subsequent increased flow rate at 100 *μl*/*min* for 1 *min* to rinse off loosely attached lipid structures.

### Giant Unilamellar Vesicle Patch SLB Composition and Formation

The lipids used for these samples are DOPC (1,2-dioleoyl-sn-glycero-3-phosphocholine, *Avanti Polar Lipids, Inc*.), Cholesterol (Ovine, cholest-5-en-3β-ol, *Avanti Polar lipids, Inc*.), Atto 488 -DOPE (1,2-Dioleoyl-sn-glycero-3-phosphoethanolamine, Atto-Tec), DSPE-peg2000-biotin (1,2-distearoyl-sn-glycero-3-phosphoethanolamine-N-[biotinyl(polyethyleneglycol)-2000] (ammonium salt), *Avanti Polar Lipids, Inc*.) and DPPE-peg2000-STAR RED (1,2-dipalmitoyl-sn-glycero-3-phosphoethanolamine-N-[azido(polyethyleneglycol)-2000, Abberior).

We prepared Giant Unilamellar Vescicles (GUVs) through electroformation similar to [20, 26] in sucrose solution (300 *mM*), using a solution of DOPC and DOPC:CHOL 7:3 depending on the experiment shown. In both cases we added DOPE Atto 488 (0.01 Mol%). In the experiments in which QDs are used we added to the lipid solution also DSPE-peg2000-Biotin (0.01 Mol%). We plasma cleaned the coverslip to rupture the GUVs and create GUV patches and we used, as buffer, phosphate buffer saline (PBS, 137*mM* NaCl, 10*mM* phosphate, 2.7*mM* KCl) to keep the supported lipid bilayer (SLB) hydrated. We then labeled the biotinylated lipids with streptavidin conjugated 655 QDs (1*μM* concentration, Invitrogen by Thermo Fisher Scientific) at a concentration of 50*pM* for MINFLUX SPT while we used streptavidin conjugated 605 QDs (1*μM* concentration, Invitrogen by Thermo Fisher Scientific) at a concentration of 1*pM* for TIRF single particle tracking. In the experiments in which we track the lipid analog DPPE-peg2000-STAR RED we labeled the SLB with a concentration of 50*pM*.

### TIRF data acquisition

The TIRF single particle tracking data are acquired on a custom made iSCAT-TIRF microscope as described in [20]. We excited the 605 QDs with a 488nm diode laser and, to efficiently detect the fluorescence signal, the filters in the TIRF’s imaging channel have been changed accordingly. We acquired 3000 frames for each measurement with an exposure time of 10*ms* that led to an effective frame time of 67*ms*.

### Iterative MINFLUX Single Particle Tracking

The SPT experiments were performed on a commercial MINFLUX microscope setup (Abberior Instruments GmbH.), which is based on an iterative localization approach [8]. For this study, we restricted our investigation to 2D tracking, although the setup is capable of 3D tracking. The device was provided with pre-determined TCP scanning parameters, that the users could modify, albeit in a limited fashion. The relevant scanning parameters used here are reported in Table 1. We note that the pattern dwell time (*dT*), i.e. the total time that the MINFLUX setup spends collecting photons during one orbit cycle, could be altered without much consequence. On the other hand, the photon limit (*PL*) and TCP diameter (*L*) required additional changes to the parameters provided for the localization correction as the CRLB (Equation 1) is dependent on both. They are available at “…/seq/containers/*.json”. We highlight that another variable changed between pattern iterations was the excitation laser power. We express this by listing amongst the parameters a laser power multiplier, which refers to a starting excitation power of 10*μW* at the sample plane.

### Iterative MINFLUX Localization Estimation

Iterative MINFLUX realizes processive particle localization by employing a chain of successive localization estimation steps. Behavior during these steps is governed by the *iMFX*-sequence parameters (Appendix 2, Parameter Overview). Each step follows the same core principles:

Around an initial assumed particle localization, the scanning beam is steered along a pre-defined path called the Target Coordinate Pattern (TCP) with its vertices on a virtual circle with diameter *L* using Electro Optical Deflectors (EODs). At each vertex, signal is integrated for a time (*t*_*integration*_) equal to *t*_*integration*_ = *t*_*dwell*_/(*N*_*vertices*_ ⋅ *N*_*TCP*_), where *t*_*dwell*_ is the Dwell Time (*dT*), *N*_*vertices*_ the number of vertices on the TCP and *N*_*TCP*_ the number of TCP *Pattern Repetitions* (*patRepeat*). Additionally, photonic signal can be integrated at a central spot to calculate the Center-Frequency-Ratio. After completing the TCP including the optional center spot, the beam is placed back in the initial position. Given its quantized and repetitive nature, we call the entire completed procession of signal integration a TCP cycle or roundtrip.

After completing a TCP, the Effective Frequency at Offset (EFO) is calculated by dividing the number of detected photons by the *dT*. If the *Automatic Background Estimation* (*bgcSense*) has been enabled, the local estimated background signal is subtracted from the EFO. The EFO is then compared against the *Background Threshold* (*bgcThreshold*) to determine if the integrated signal can be considered valid.

If the EFO is below the Background Threshold, another roundtrip is kicked off and the collected photons discarded.

If the EFO surpasses the *Background Threshold*, the collected photons are counted against the Photon Limit (*PL*), i.e. the minimal number of photons requested to be used for a localization estimation.

Should the number of detected photons above the background threshold collected be less than the *PL*, another TCP cycle is started and the photons retained.

Should the number of valid photons collected be equal or surpass the *PL*, the MINFLUX microscope will start to calculate a next localization estimate, which after being corrected in a second step using the *Estimation Coefficients* (*estCoefficients*) is used as the initial particle position for the next run. A Galvanometric Scanner is used to re-center the virtual circle around that position and routine is re-engaged.

Our experimental hardware highlights that moving the donut beam center between vertices takes about *t*_*EOD*_ ≈ 5*μs*, while the localization estimation and TCP repositioning takes about *t*_*REP*_ ≈ 20*μs*. This enables us to write down an ideal time-to-localization *t*^2*D*^ for a single *iMFX* determined particle localization in 2D:

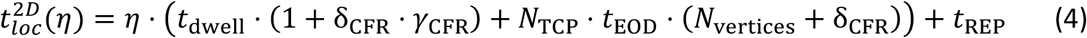

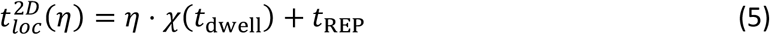

Where *η* ∈ [1, ∞) is the number of roundtrips completed, θ_CFR_ is a delta-function that returns δ_CFR_ = 1 if the CFR-Check is enabled and *γ*_CFR_ is a factor that determines the amount of time spent integrating the signal for the *CFR-Check* (*ctrDwellFactor*). We summarize the constant part of Equation 4 into *χ*(*t*_dwell_) in Equation 5.

To display the split between signal integration 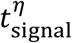 and hardware overhead 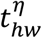, we can express Equation 4 in the following way:

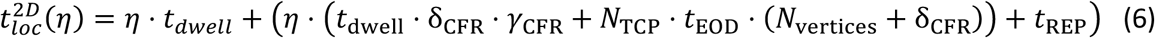

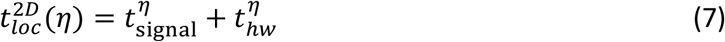

### Temporal Optimization

Considering linear Equation 5, we determine two angles of minimization, the number of TCP round trips *η* and the slope *χ*(*t*_dwell_). While minimizing the latter is trivial considering Equation 4, we must understand *η* as a function of the laser-power, *PL*, and *dT* that minimizes exactly when at least *PL* number of photons arrive at the detector within *dT* seconds. Providing a complete mathematical model of this behavior is beyond the scope of this here work, but for our purposes it is sufficient to understand that these three factors, i.e. laser-power, *PL*, and *dT*, play an essential role when trying to optimize *iMFX*-enabled SPT experiments.

### Turning Angle Distribution

Turning angles are defined as the angle between the displacement vectors 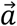 and 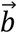 which describe the particle motion along three consecutive positions A, B, C. It is calculated as follows:

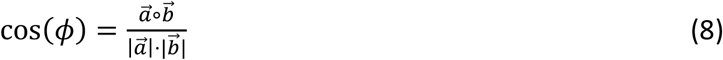

The Turning Angle Distribution (TAD) can be used to investigate directionality between successive datapoints, hence posing as a tool to describe single particle motion across time.

### Number of Cycles

Assuming a steady fluorescent signal, we can straightforwardly extract the number of cycles *η* required for each localization in *iMFX* from the data provided as follows:

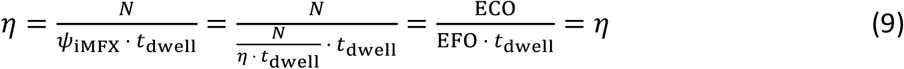

Where *N* (ECO) is the number of photons acquired at the TCP during position estimation, *ψ*_iMFX_ (EFO) is the photon detection frequency at the TCP and *t*_dwell_ is the dwell time as put in the sequence.

### Ergodic Hypothesis

The ergodic hypothesis, the only fundamental assumption in equilibrium statistical physics [27], implies that the average of a process parameter over time would equate the average over the complete statistical ensemble. In the case of particle diffusion, this relates to:

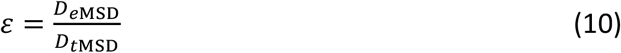

Where *D*_*t*MSD_ is the time-average diffusion coefficient and *D*_*e*MSD_ is the ensemble-average diffusion coefficient.

### Defining an Upper Limit for Trackable Diffusion Rates

Assuming Brownian particles to exhibit a constant average photon emission rate ⟨θ_p_⟩ and the average photon detection rate ⟨*ψ*_iMFX_⟩ ≤ ⟨θ_p_⟩, we can give an estimation of the upper limit of possibly trackable diffusion rates 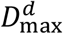 given 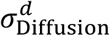 and that the particle needs to remain within the area of the TCP during the localization process.

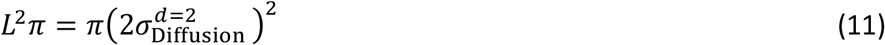

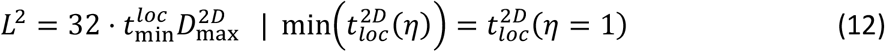

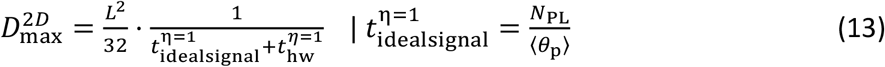

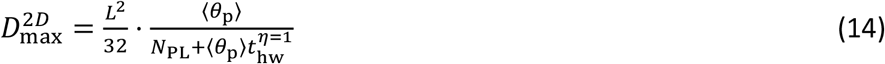

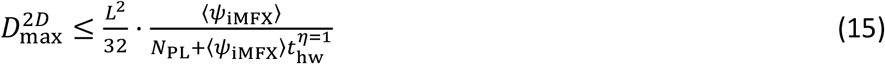

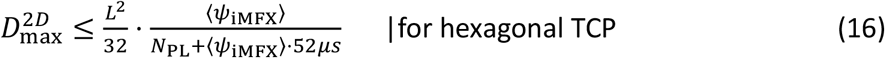

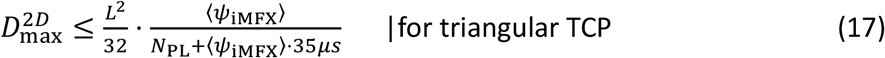

## Supporting information

Appendix I - Optimization Guide

Appendix II - Parameter Overview

## Code Availability

Due to it being highly relevant to ongoing projects, we decided to postpone releasing our custom code on GitHub to a later date. However, any code will be made available to both editors and reviewers upon request.

## Data and Sequence Availability

Any data and all sequences used in the manuscript of this current stage are made available withing the following GitHub repository. They are named based on their respective experimental parameters. Further, we include a toolbox to conveniently access the data.

## Acknowledgements

The authors greatly acknowledge financial support by the Deutsche Forschungsgemeinschaft (DFG, German Research Foundation; Instrument funding MINFLUX Jena INST 275_405_1; Germany’s Excellence Strategy – EXC 2051 – Project-ID 390713860; project number 316213987 – SFB 1278; GRK M-M-M: GRK 2723/1 – 2023 – ID 44711651), the State of Thuringia (TMWWDG), the Leibniz Association (Leibniz ScienceCampus InfectoOptics Jena financed by the funding line Strategic Networking of the Leibniz Association, project number W8/2018; and Leibniz Collaborative Excellence Programme, project AMPel – project numer K548/2023), the Free State of Thuringia (TAB; Advanced Flu-Spec / 2020 FGZ: FGI 0031), and the Alexander von Humboldt Foundation (Research Group Linkage Fund). Further, this work is supported by the BMBF, funding program LIVE2QMIC (FGZ: 13N15956) as well as Photonics Research Germany (FKZ: 13N15713 / 13N15717) and is integrated into the Leibniz Center for Photonics in Infection Research (LPI). The LPI initiated by Leibniz-IPHT, Leibniz-HKI, UKJ and FSU Jena is part of the BMBF national roadmap for research infrastructures.

## References

[1] Manzo C, Garcia-Parajo MF. A review of progress in single particle tracking: from methods to biophysical insights. Rep Prog Phys 2015; 78(12): 124601 [10.1088/0034-4885/78/12/124601][PMID: 26511974]

[2] Jacobson K, Liu P, Lagerholm BC. The Lateral Organization and Mobility of Plasma Membrane Components. Cell 2019; 177(4): 806–19 [10.1016/j.cell.2019.04.018][PMID: 31051105]

[3] Qian H, Sheetz MP, Elson EL. Single particle tracking. Analysis of diffusion and flow in two-dimensional systems. Biophys J 1991; 60(4): 910–21 [10.1016/S0006-3495(91)82125-7][PMID: 1742458]

[4] Gurtovenko AA, Javanainen M, Lolicato F, Vattulainen I. The Devil Is in the Details: What Do We Really Track in Single-Particle Tracking Experiments of Diffusion in Biological Membranes? J Phys Chem Lett 2019; 10(5): 1005–11 [10.1021/acs.jpclett.9b00065][PMID: 30768280]

[5] Wieser S, Schütz GJ. Tracking single molecules in the live cell plasma membrane-Do’s and Don’t’s. Methods 2008; 46(2): 131–40 [10.1016/j.ymeth.2008.06.010][PMID: 18634880]

[6] Lanzanò L, Gratton E. Orbital Single Particle Tracking on a commercial confocal microscope using piezoelectric stage feedback. Methods Appl. Fluoresc. 2014; 2(2): 24010 [10.1088/2050-6120/2/2/024010][PMID: 25419461]

[7] Balzarotti F, Eilers Y, Gwosch KC, et al. Nanometer resolution imaging and tracking of fluorescent molecules with minimal photon fluxes. Science 2017; 355(6325): 606–12 [10.1126/science.aak9913][PMID: 28008086]

[8] Schmidt R, Weihs T, Wurm CA, et al. MINFLUX nanometer-scale 3D imaging and microsecond-range tracking on a common fluorescence microscope. Nat Commun 2021; 12(1): 1478 [10.1038/s41467-021-21652-z][PMID: 33674570]

[9] Einstein A. Über die von der molekularkinetischen Theorie der Wärme geforderte Bewegung von in ruhenden Flüssigkeiten suspendierten Teilchen. Annalen der Physik 1905; 322(8): 549–60 [10.1002/andp.19053220806]

[10] Aberior Instruments GmbH. Personal Communication with Abberior Instruments GmbH. Hans-Adolf-Krebs-Weg 1, Weende, 37077 Göttingen, Germany; 2024 2024 Jul 16.

[11] Aberior Instruments GmbH. Personal Communication with Abberior Instruments GmbH; 2024 2024 Nov 6.

[12] Tabaei SR, Choi J-H, Haw Zan G, Zhdanov VP, Cho N-J. Solvent-assisted lipid bilayer formation on silicon dioxide and gold. Langmuir 2014; 30(34): 10363–73 [10.1021/la501534f][PMID: 25111254]

[13] Invitrogen TFS. Qdot® Streptavidin Conjugates; 2007 [cited 2025 May 15] Available from: URL: https://assets.thermofisher.com/TFS-Assets/LSG/manuals/mp19000.pdf.

[14] Reina F. Applications of interferometric scattering microscopy (iSCAT) to single particle tracking in model and cell membranes. University of Oxford.

[15] Mascalchi P, Haanappel E, Carayon K, Mazères S, Salomé L. Probing the influence of the particle in Single Particle Tracking measurements of lipid diffusion. Soft Matter 2012; 8(16): 4462 [10.1039/c2sm07018a]

[16] Clausen MP, Lagerholm BC. Visualization of plasma membrane compartmentalization by high-speed quantum dot tracking. Nano Lett 2013; 13(6): 2332–7 [10.1021/nl303151f][PMID: 23647479]

[17] Clausen MP, Lagerholm BC. The probe rules in single particle tracking. Curr Protein Pept Sci 2011; 12(8): 699–713 [10.2174/138920311798841672][PMID: 22044141]

[18] Vogler BTL, Reina F, Eggeling C. Blob-B-Gone: a lightweight framework for removing blob artifacts from 2D/3D MINFLUX single-particle tracking data. Front Bioinform 2023; 3:p 1268899 [10.3389/fbinf.2023.1268899][PMID: 38076029]

[19] Michalet X, Berglund AJ. Optimal diffusion coefficient estimation in single-particle tracking. Phys Rev E Stat Nonlin Soft Matter Phys 2012; 85(6 Pt 1): 61916 [10.1103/PhysRevE.85.061916][PMID: 23005136]

[20] G. De Angelis, J. Abramo, M. Miasnikova, M. Taubert, C. Eggeling, and F. Reina. Homogeneous large field-of-view and compact iSCAT-TIRF setup for dynamic single molecule measurements. Optics Express. 2024 Dec 10; 32(26):46607–20 2024.

[21] Spillane KM, Ortega-Arroyo J, Wit G de, et al. High-speed single-particle tracking of GM1 in model membranes reveals anomalous diffusion due to interleaflet coupling and molecular pinning. Nano Lett 2014; 14(9): 5390–7 [10.1021/nl502536u][PMID: 25133992]

[22] Mobarak E, Javanainen M, Kulig W, et al. How to minimize dye-induced perturbations while studying biomembrane structure and dynamics: PEG linkers as a rational alternative. Biochim Biophys Acta Biomembr 2018; 1860(11): 2436–45 [10.1016/j.bbamem.2018.07.003][PMID: 30028957]

[23] Honigmann A, Mueller V, Hell SW, Eggeling C. STED microscopy detects and quantifies liquid phase separation in lipid membranes using a new far-red emitting fluorescent phosphoglycerolipid analogue. Faraday Discuss 2013; 161: 77-89; discussion 113-50 [10.1039/c2fd20107k][PMID: 23805739]

[24] Wirth JO, Schentarra E-M, Scheiderer L, Macarrón-Palacios V, Tarnawski M, Hell SW. Uncovering kinesin dynamics in neurites with MINFLUX. Commun Biol 2024; 7(1): 661 [10.1038/s42003-024-06358-4][PMID: 38811803]

[25] Deguchi T, Iwanski MK, Schentarra E-M, et al. Direct observation of motor protein stepping in living cells using MINFLUX. Science 2023; 379(6636): 1010–5 [10.1126/science.ade2676][PMID: 36893247]

[26] Méléard P, Bagatolli LA, Pott T. Giant unilamellar vesicle electroformation from lipid mixtures to native membranes under physiological conditions. Methods Enzymol 2009; 465:p 161–76 [10.1016/S0076-6879(09)65009-6][PMID: 19913167]

[27] Wen B, Li M-G, Liu J, Bao J-D. Ergodic Measure and Potential Control of Anomalous Diffusion. Entropy (Basel) 2023; 25(7) [10.3390/e25071012][PMID: 37509959]

