## Appendix I - Optimization Guide for "Parameter Optimization for Iterative MINFLUX Microscopy enabled Single Particle Tracking"

### Iterative-MINFLUX Sequence Optimization Guide for 2D SPT Experiments

#### Step 1 Preliminary Questions

Ask yourself the following questions:

- What exactly is the phenomenon of interest?
  - What are the relevant spatial and temporal scales?
  - What is the lateral localization precision necessary to observe it?
- Which is the priority, sampling rate or localization precision?
- Is this an absolute measurement or do I just want to compare different experimental conditions?
- Is the effect I want to observe faster than $50\mu s$?

**Note**: If the required sampling rate is faster than $20kHz$ ($50\mu s$ localization time), a different approach than MINFLUX is necessary.

#### Step 2 Fluorescent Label and Optimization of Fluorescence Detection

The goal of this step is to adjust the excitation intensity such that enough photons can be detected in as little time as possible while avoiding photo-damage to the sample as much as possible. A handy guideline is to make sure that the average photon detection rate from the sample, given a known Dwell Time (*dT*) and Photon Limit (*PL*), is sufficient to obtain the whole photon budget in one iteration. For example, for $PL = 10$ and $dT = 100\mu s$, the target average photon detection rate is at least 100kHz, that is, 10 photons per $100\mu s$.

The size of the emitting particle must be kept in mind as larger targets, e.g. quantum dots, may not be approximated as a point coordinate as the lateral resolution of the microscope is in the same order of magnitude as the spatial extent of the tracked object, which introduces a possible further source of localization error to the measurement.

**TIP**: Measuring the average photon detection rate of your target at different excitation laser powers may be useful, to evaluate, for example, the influence of photophysical effects such as blinking or flickering of the fluorescent dyes.

***Warning***: MINFLUX requires extremely low concentrations of target molecules to ensure tracking and reduce the likelihood of mid-trace-target-swap events.

### Step 3 Adjust the MINFLUX Sequence

#### How are Iterative-MINFLUX Measurements Initiated and Controlled?

Once a suitable ROI is selected in the sample, a MINFLUX measurement can be initiated, and the device will follow a *sequence* containing the instructions to perform it. A MINFLUX sequence is a *.json* file that dictates how the microscope hardware operates once the measurement is started. As the localization routine operates in sequential iterations in *iMFX*, there are two types of parameters involved: global ones affecting the entire measurement and local ones affecting only their respective iteration.

#### General Advice for Modifying Iterative-MINFLUX Sequence Parameters

When changing the laser power and/or Power Factor parameters, it may be necessary to re-evaluate the background threshold. The same is true when changing the setting for the Automatic Background Estimation (given as *bgcSense* in the sequence file), which can prove useful for highly heterogeneous samples.

When changing *PL* and TCP diameter size (given as *patGeoFactor* in the sequence file), the correct estimation coefficients need to be entered in the sequence files. These are usually provided by the manufacturer (Abberior Instruments GmbH) and can be easily copied into the sequence at the relevant place. If precise estimation coefficients do not exist for the chosen combination of *PL* and *patGeoFactor*, it is possible to choose the next closest combination if the deviation is minimal or reach out to the manufacturer to seek further advice.

#### Recommendations for Global Parameters

- Disable damping [*damping*: 0] to follow the particle more closely. If severe overshoot is present in the final datasets, or if the measurement is not intended to measure absolute quantities (such as diffusion rates), it is possible to set [*damping*: 1] to avoid possible mechanical overshoots in the scanning hardware when following the particle. It is, however, not recommended to increase it beyond 1, which would result in worse tracking performance.
- Make sure to only repeat the last iteration when tracking by setting [*headstart*: -1]. This is essential to ensure fast and reliable tracking.
- Enable permissive tracking [*stickiness*: $> 0$] to prevent the routine from being disrupted prematurely due to target particle flickering or blinking events. While this may introduce mid-trace-target-swap events it greatly increases the continuity of SPT process. Additionally, these events can be filtered out in post processing with relative ease, given that they introduce a significant temporal gap when they occur.
- (Optional) Turn on the Automatic Background Estimation [*bgcSense*: true] for cell samples. This will enable the MINFLUX to gauge the local background level for each position in the searching phase of the sequence. This is essential for samples with spatially inhomogeneous background.

**TIP**: Use the *locLimit* parameter to set a maximum number of consecutive localizations per continuous trace. This has two major benefits: first, it will increase the number of different particles tracked by forcing the routine to return to search mode after the localization limit has been reached, and also it reduces the chance of the tracking routine getting "stuck" on of immobile particles that may be present in the sample.

#### Choosing the TCP size *L* of the Final Iteration

- Fast diffusion, e.g. on SLBs -> L = 150nm [*patGeoFactor*: 0.42]
- Medium-Slow diffusion, e.g. on cells -> L = 100nm [*patGeoFactor*: 0.28]
- Slow diffusion, e.g., molecular motors -> L = 75nm [*patGeoFactor*: 0.21]

Generally speaking, the faster the particle, the larger the L should be. A smaller TCP diameter is especially recommended if optimizing for spatial precision.

#### Choosing the TCP Geometry for the Final Iteration

- Generally, or in doubt -> Hexagonal pattern
- Only in specific cases, e.g. unusually dim target -> Triangular pattern

It is strongly recommended to always use the hexagonal pattern for tracking as it offers, in principle, superior precision compared to the triangular pattern, in terms of the symmetry of the precision of localization throughout the TCP area. However, restricting the pattern to three vertices increases the average signal integration time per vertex, i.e., TCP scanning position, which may be necessary for fast moving particles with dim fluorescence targets. Due to the reduced number of vertices, a triangular scanning pattern takes about $15\mu s$ less to complete compared to the hexagonal one.

#### Adjust the Background Threshold

- Adjust your sequence to measure the background by setting the background threshold and photon limit to zero [*bgcThreshold*: 0, *phtLimit*: 0] or pick one of our example sequences.
- Conduct a MINFLUX measurement and estimate a background threshold for the final iteration from the average of the EFO histogram obtained from the *paraFLUX* software provided by Abberior instruments GmbH.

**Tip**: Do not forget to reset the background threshold and photon limit for all iterations should you want to use the same sequence for tracking later.

#### Step 4 Engineer the Final Sequence

##### Initial Conditions

It is recommended to start with a sequence where the final iteration has a permissive photon limit, i.e. with initial parameters that provide fast and reliable sampling with a small time to localization with less emphasis on minimizing the localization error, for example:

- Photon Limit: 10 [*phtLimit*: 10]
- Dwell Time: $100\mu s$ [*patDwellTime*: 100e-6]
- Power Factor: 3 [*pwrFactor*: 3.0]

With such parameters, it is possible to collect the necessary data to further optimize the MINFLUX sequence and tailor it to the experiment.

**Tip**: Using the Power Factor parameter, it is possible to control the laser power to use during the final iteration. This applies exclusively during the iteration in which it is included, providing the necessary excitation power to produce the desired photon detection rate exclusively during the tracking step, and not before.

##### Generally recommended settings for MINFLUX SPT experiments

- Disable preset TCP pattern repetitions [*patRepeat*: 1] to reduce the overall average time-to-localization $\langle t_{loc} \rangle$ as each additional repetition adds significant hardware time overhead.
- Disable the Center-Frequency-Ratio Check [*ccrLimit*: -1.0] in the last iteration in the sequence to prevent additional time overhead and significantly reduce the time-to-localization and photonic-stress to the sample by removing an additional detection at the center of the TCP.

##### Optimization Criteria

Using the thus prepared sequence, take an initial MINFLUX dataset of the labelled sample. Evaluate your data using the tools provided here or your own and further optimize the sequence until ideal conditions are reached. We recommend determining and using the following values to optimize the sequence, whose definition are present in the main text:

Acquisition Speed:

- $\langle \eta \rangle$ – The average number of cycles per localization (Use when adjusting PL)
  To ensure fast and homogeneous tracking, optimize $\langle \eta \rangle = \frac{\langle N \rangle}{\langle \psi_{\mathrm{iMFX}} \rangle \cdot t_{dwell}} = 1.$
- $\langle t_{loc} \rangle$ – The average time-to-localization (Use when adjusting dT)
  To speed up tracking and catch faster particles, optimize $\langle t_{loc} \rangle \rightarrow 0$

Acquisition Fidelity and Photonic Stress:

- $\frac{\langle N \rangle}{\langle \eta \rangle \cdot N_{PL}}$ – Multiples of the photon limit detected per localization
  To ensure fast and homogeneous tracking, optimize $\frac{\langle N \rangle}{\langle \eta \rangle \cdot N_{PL}} > 1$

Where $\langle\psi_{\mathrm{iMFX}}\rangle$ is the average photon detection frequency per localization sometimes referred to as EFO, $\langle N\rangle$ $\left(= \langle\psi_{\mathrm{iMFX}} \cdot t_{\mathrm{signal}}^{\eta}\rangle = \langle\psi_{\mathrm{iMFX}} \cdot \eta \cdot t_{\mathrm{dwell}}\rangle = \langle\psi_{\mathrm{iMFX}}\rangle \cdot \langle\eta\rangle \cdot t_{\mathrm{dwell}}\right)$ the average number of collected photons sometimes referred to as ECO, $t_{\mathrm{loc}} = t_{\mathrm{signal}}^{\eta} + t_{hw}^{\eta}$ (Equations 4-7, main text) the total time needed for a localization with number of cycles $\eta$, $t_{\mathrm{dwell}}$ the dwell time per localization round as put in the sequence, and $N_{PL}$ the photon limit.

Target Values:

It is useful to have some reference values to compare the results of the MINFLUX experiment. In the case of diffusion measurements, this would be an initial estimate for the diffusion rate $\langle \mathrm{D}_{\mathrm{MSD}}\rangle$ taken from previous experiment and/or for simulations.

**Tip**: It is recommended to examine the EFO histogram during the measurement as follows: The number of Poissonian modes (Optimization Guide Figure 1) provides an approximation of the average number of cycles per localization $\langle\eta\rangle$, which itself provides an estimate of $\langle t_{loc}\rangle$. The variance of the 1-cycle-distribution (See red line in Optimization Guide Figure 1) may be used to gauge photon-use-efficiency $\frac{\langle N\rangle}{\langle\eta\rangle \cdot N_{PL}}$.

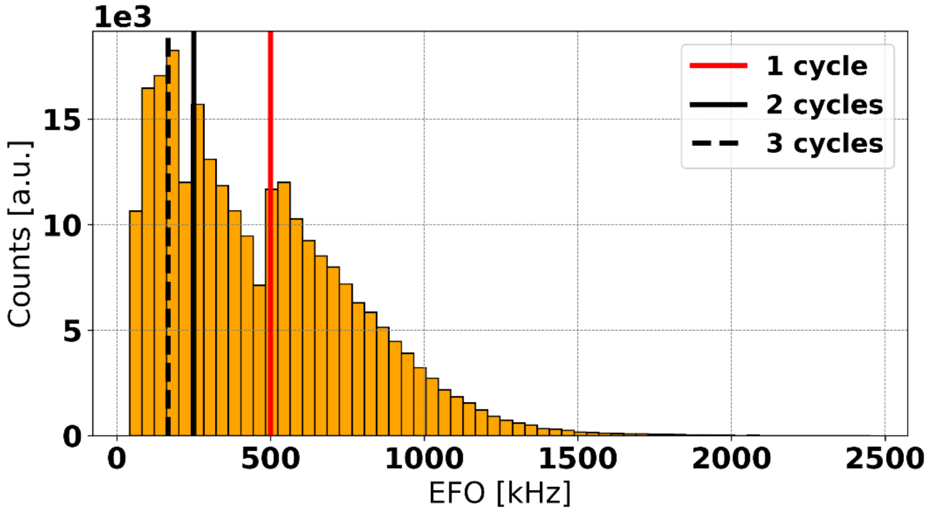


***Optimization Guide Figure 1 -*** Exemplary EFO histogram for the Brownian diffusion of a fluorescent lipid analogue on a GUV-patch SLB (see Materials and Methods) tracked using Iterative-MINFLUX. Main sequence parameters are PL 50, dT100 and DMP0 (See Main Text, Table 1). Horizontal lines have been added to visualize the difference between the different number of cycles $\eta$. The average number of cycles per localization for this dataset is $\langle\eta\rangle = 2$ (See Main Text, Table 2).

#### Step 5 Optimization Loop

1. **Perform a MINFLUX experiment and calculate $\langle \boldsymbol{\eta} \rangle$ and $\langle \boldsymbol{t_{loc}} \rangle$.**
2. **If $\langle \boldsymbol{\eta} \rangle = \mathbf{1}$**: *Advance to 3*.
   **If $\langle \boldsymbol{\eta} \rangle > \mathbf{1}$**: MINFLUX tracking is too slow. Try decreasing the PL or increasing the Power Factor if the sample and background level permit it. *Adjust and start over*.
3. **If $\langle \boldsymbol{t_{loc}} \rangle \leq \boldsymbol{t_{trg}}/\mathbf{2}$** (with $t_{trg}$ the timescale of the effect under study): *Advance to 4*.
   **If $\langle \boldsymbol{t_{loc}} \rangle > \boldsymbol{t_{trg}}/\mathbf{2}$**: Try lowering the *dT*. Alternatively, although we do not recommend it, consider changing to the triangular TCP pattern which is marginally faster than the hexagonal. *Adjust and start over*.
4. **If $\mathbf{1} < \frac{\langle \boldsymbol{N} \rangle}{\langle \boldsymbol{\eta} \rangle \cdot \boldsymbol{N_{PL}}} < \mathbf{1.5}$**: every photon acquired per cycle is used effectively.
   *Advance to 5*.
   **If $\frac{\langle \boldsymbol{N} \rangle}{\langle \boldsymbol{\eta} \rangle \cdot \boldsymbol{N_{PL}}} > \mathbf{1.5}$**: there are photons to spare. Think about increasing the PL or decreasing *dT* to increase the localization precision or tracking speed or consider lowering the Power Factor to decrease photonic stress and potentially increase the track length. *Adjust and start over*.
5. **If reference experiments exist**:
   *Calculate and evaluate $\langle D_{MSD} \rangle$:*
      *If $\langle D_{MSD} \rangle$ matches the expectation, this concludes this optimization scheme*.
      *If not, it might be necessary to re-evaluate the results of 2. and 3.*
   **If no reference experiments exist**:
      *This concludes this optimization scheme.*
   **Tip**: Whether there are reference experiments or not, it is almost always possible to optimize for the localization precision by using the dynamic localization error $\langle \sigma_{\mathrm{MSD}} \rangle$ as target value.

With this optimization loop we have three goals: ensuring fast, homogeneous and reliable tracking by measuring with a single exposure cycle per localization ($\langle \eta \rangle = 1$), and to match the timescale of the experiment as closely as possible while satisfying Nyquist sampling requirements ($\langle t_{loc} \rangle \leq t_{trg}/2$) all while trying to use the photons at our disposal as efficiently as possible increasing the localization precision and reducing photonic stress on the sample $\left(1 < \frac{\langle N \rangle}{\langle \eta \rangle \cdot N_{PL}} < 1.5\right)$.

Tip: If the detected trajectories are extremely short overall, the reason could be that the excitation laser power or Power Factor are too high, bleaching the target very fast, or there could be experiencing focus drift. If the final data set is split between extremely long and extremely short traces, it is more than likely that the MINFLUX sequence is too slow, in which case the results will be heavily biased towards slow and/or immobile particles.

#### Adjustment Limits

The relevant parameters for optimization of SPT measurements in MINFLUX can be adjusted within the limits below (Optimization Guide Table 1), although it must be kept in mind that any adjustment still needs to be allowed by the set of localization estimation coefficient provided by the manufacturer of the setup.

**Optimization Guide Table 1** – Suggestions for the lower and upper limit for the Photon Limit, Dwell Time, and Power Factor *iMFX* parameters.

| Parameter | Lower Limit | Upper Limit |
|---|---|---|
| **Photon Limit** | Equal to the number of vertices of the TCP pattern. | No upper limit. |
| **Dwell Time** | Equal to the hardware overhead $t_{hw}^{\eta=1}$ ($52\mu s$ for the hexagonal scanning pattern and $35\mu s$ for the triangular pattern). | Half of the relevant timescale of the phenomenon of interest in the experiment, to satisfy Nyquist sampling conditions. |
| **Power Factor** | 1.0, equal to the excitation power input for the experiment. | No upper limit, but photobleaching and other photophysical effects must be kept in mind. |

#### A General Rule of Thumb

The laser power and *PL* should be kept as high as possible and the *dT* as low as possible while maintaining $\langle\eta\rangle = 1$. That way the time-to-localization is automatically minimized, while maximizing lateral localization precision and tracking rate.
