## Appendix II - Parameter Overview for "Parameter Optimization for Iterative MINFLUX Microscopy enabled Single Particle Tracking"

### Overview of the Most Influential Sequence Parameters in Iterative-MINFLUX-enabled Single Particle Tracking

#### Disclaimer:

**The following list of parameters, parameter names, and descriptions given below is subject to change. Any information is provided as is for the MINFLUX-*iMSPECTOR* version *v16.3.15635* supplied by the manufacturer (Abberior Instruments GmbH).**

#### Global Parameters

##### Automatic Background Estimation

**Parameter Name:** *bgcSense*

**Explanation:** Determines if and how many times the background level is gauged during grid search. A nonzero (or true) value indicates that background measurements are performed. When an integer $n > 1$ is provided as input, the background is estimated in increments of $n$, i.e. at each $n$-th time that no particle is found within a grid cell, while a value of zero (or false) turns background sensing off.

The background level estimate will be subtracted from the signal obtained in each iteration before the background threshold is applied.

**Tip:** This is advantageous for heterogeneous samples, e.g. cellular membranes. The Background Threshold should be fit to the sample whether the automatic estimation is used or not. Make sure to use the same value, i.e. the same setting, for *bgcSense* when adjusting the Background Threshold for the sample used.

**Expected Input:** Integer (typical values within 0–10) or Boolean.

##### Center Dwell Factor

**Parameter Name:** *ctrDwellFactor*

**Explanation:** Controls the fraction of the total dwell time (see *patDwellTime* below) allocated for a CFR-Check. When active, the CFR-Check duration is computed as:

$$t_{center} = ctrDwellFactor \cdot patDwellTime$$

**Tip:** Keep in mind that the set value determines the time spent in the center spot regardless of the iteration.

**Expected Input:** Positive Floating-Point Value.

#### Damping

**Parameter Name:** *damping*

**Explanation:** Artificially shortens the update distance between consecutive localizations by a factor:

$$\Delta_{update} = \Delta_{real} \cdot 2^{-damping}$$

**Tip:** Damping may be used to counter localization overshoot or to mitigate possible jitters introduced by the galvanometric scanner. However, it is discouraged to use during Single Particle Tracking (SPT) as it arbitrarily modifies the results underestimating particle motility (Supplementary Figure 1).

**Expected Input:** Positive Floating-Point Value.

#### Localization Limit per continuous Trace

**Parameter Name:** *locLimit*

**Explanation:** Sets an upper threshold for the number of valid consecutive localizations allowed along a single trace. The SPT process is stopped after reaching the threshold and returns to the grid search.

**Expected Input:** Positive Integer or –1 (to disable the limit).

#### Number of Permitted Localization Attempts

**Parameter Name:** *stickiness*

**Explanation:** Defines the number of attempts made to localize a particle. An attempt is terminated by a failed CFR-Check or a break condition triggered by exceeding the *maxOffTime*.

**Tip:** During SPT it is advantageous to use a permissive sequence that doesn't terminate immediately to improve tracking consistency. Keep in mind that this, however, requires post processing to get rid of mid-trace-particle-swap events.

**Expected Input:** Positive Integer (with 0 disabling the check).

#### Sequence Name

**Parameter Name:** *id*

**Explanation:** Set a name or identifier for the sequence.

**Tip:** This is the name that will appear in the *iMSPECTOR* software during sequence selection. It is encouraged to use the filename of the sequence *.json* file for easy readability.

**Expected Input:** String.

#### Sequence Repetition Entry Point

**Parameter Name:** *headstart*

**Explanation:** Specifies from where to re-enter the sequence after the final iteration has concluded with a particle localization estimated. Positive numbers start counting from the top, negative from the bottom.

**Tip:** As we aim for speed and photon efficiency during SPT it is highly encouraged to repeat only the final iteration ($headstart = -1$).

**Expected Input:** Integer.

#### Local Parameters

##### Background Threshold

**Parameter Name:** *bgcThreshold*

**Explanation:** Dictates the baseline threshold (in Hz) for the background level as a high-pass filter. After each roundtrip, the system determines if the obtained signal can be considered valid foreground. Signal frequencies beyond this value will be considered valid to use for the localization estimation. In case the signal resides below the threshold, another roundtrip is engaged.

**Tip:** Make sure to adjust the threshold to on a per-experiment basis. Think about the expected sample homogeneity and consider whether to additionally use *Automatic Background Estimation*. Make sure however to stick with the decision and adjust the thresholds of the sequence according to the settings used during tracking.

**Expected Input:** Positive Integer or –1 (to disable).

##### Center-Frequency-Ratio Check (CFR Check)

**Parameter Name:** *CCRLimit*

**Explanation:** This activates the CFR-Check in the respective iteration when set to a positive value. This means that after finishing the TCP roundtrip, the beam will be placed in the center of the pattern and signal will be integrated for a time dictated by the Center Dwell Factor. After that the Effective Frequency of the Center (EFC) will be compared to the Effective Frequency at the Offset (EFO), i.e. the signal obtained at the vertices, to calculate the CFR:

$$\mathrm{CFR} = \mathrm{EFC} / \mathrm{EFO}$$

If the ratio exceeds the value set for the *CCRLimit*, the track is terminated, and an attempt is concluded.

**Tip:** It is worth noting that due to the small TCP diameter and the fluorescent target constantly moving, the CFR-Check is not a suitable measure to be used during the tracking of free and or fast particles and should hence be turned off in the final iteration. Additionally, deactivation of the CFR-Check significantly speeds up data acquisition and thus the entire tracking routine. In case the target particle is expected to exhibit sufficiently slow and/or semi-static movement and when sampling speed is not of the essence, it could however be worth considering employing the CFR-Check in the final iteration.

**Expected Input:** Positive Floating-Point Number or –1 (to disable).

#### Dwell Time

**Parameter Name:** *patDwellTime*

**Explanation:** Specifies the time in seconds the system spends on signal integration on a TCP roundtrip. For each vertex (excluding the extra center spot if the CFR-Check is enabled), the integration time $t_{\text{integrate}}$ is given by:

$$t_{\text{integrate}} = patDwellTime \, / \, N_{\text{vertices}}$$

where $N_{vertices}$ is the number of vertices of the TCP.

**Tip:** Even in ideal conditions (i.e. $\langle\eta\rangle = 1$) the Dwell Time does not equate the time-to-localization as there is hardware induces temporal overhead $t_{hw}^{\eta=1}$ (compare Equation 7, main text) of about $t_{\text{HEX}}^{\eta=1} \approx 52\mu s$ (Hexagonal Pattern) or $t_{\text{TRI}}^{\eta=1} \approx 35\mu s$ respectively. It is discouraged to choose a Dwell Time that approaches the hardware overhead or goes below it.

**Expected Input:** Positive Floating-Point Number.

#### Estimation Coefficients

**Parameter Name:** *estCoeff*

**Explanation:** These coefficients provided by the manufacturer (Abberior Instruments GmbH) are used during the localization estimation and correct an initial estimate based on an expected number of photons and a given TCP shape and diameter.

**Tip:** Make sure to update these coefficients any time the Photon Limit, TCP Diameter and/or TCP Pattern are changed. Some coefficient configurations can be found in the premade iteration blocks called *containers* or be provided by the manufacturer (Abberior Instruments GmbH) upon request. It is possible to vary the Photon Limit slightly for an iteration keeping the same Estimation Coefficients without much decrease in localization performance, though it is encouraged to always use the correct Estimation Coefficients for each iteration to prevent errors and unwanted side effects.

**Expected Input:** A list of Floating-Point Numbers.

#### Laser Power Factor

**Parameter Name:** *pwrFactor*

**Explanation:** Adjusts the excitation laser power used in the respective iterations by multiplicatively scaling the baseline laser power by the set value.

**Tip:** It can be used to increase the laser power exclusively during tracking while keeping the photonic stress to a minimum during the rest of the experiment. However, it is advised to keep variation in this value to a minimum and rather adjust the baseline laser power specified in the *iMSPECTOR* software.

**Expected Input:** Positive Floating-Point Number.

#### Pattern Repeat

**Parameter Name:** *patRepeat*
**Explanation:** Specifies the number of times the TCP is repeated, during a single Dwell Time. The overall sum of the integration time spent at the vertices on all roundtrips is equal to the Dwell Time:

$$patDwellTime = \sum_i t^i_{integrate} = \sum_i patDwellTime \,/\, (N_{\text{vertices}} \cdot patRepeat)$$

**Tip:** Increasing the repeat count can help counteract issues such as fluorophore blinking or instrument vibrations. Keep in mind however that additional roundtrips add additional hardware overhead to the total time-to-localization (see Dwell Time).
**Expected Input:** Positive Integer (0 breaks the sequence as no scan is conducted)

#### Photon Limit

**Parameter Name:** *phtLimit*
**Explanation:** Specifies the minimum number of photons required for a valid localization estimation. If not reached within a single TCP roundtrip, another is engaged, provided no other break criterium is triggered (see Stickiness, CFR-Check and Single Interval Linger Time as well as the Methods part of the Main document).
**Tip:** For larger changes, it is recommended to use the premade iteration blocks taken from the container sequences.
**Expected Input:** Positive Integer.

#### TCP Diameter L

**Parameter Name:** *patGeoFactor*
**Explanation:** Controls the diameter of the TCP by scaling the nominal value ($360nm$). The TCP diameter calculates as follows:

$$L = patGeoFactor \cdot 360nm \cdot (wavelength/642nm)$$

Where *wavelength* is an additional iteration specific reference parameter found in the sequence but excluded in our overview that should not be edited whatsoever.
**Tip:** It is recommended to use the premade blocks of parameters called *containers* whenever changing the TCP diameter or contacting the manufacturer (Abberior Instruments GmbH).
**Expected Input:** Positive Floating-Point Number.

#### TCP Pattern

**Parameter Name:** *pattern*
**Explanation:** Defines the TCP scan pattern by selecting one of several preset geometries (such as “hexagon”, “square”, or “triangle”). This directly influences the number of vertices and the overall photon collection geometry.
**Tip:** It is highly suggested not to change the pattern but to rely on the premade iteration blocks called *containers* provided by the manufacturer (Abberior Instruments GmbH).
**Expected Input:** String (referencing a preset pattern).

#### Single Interval Linger Time

**Parameter Name:** *maxOffTime*

**Explanation:** Determines a grace interval in seconds that the signal can remain below the Background Threshold during signal integration. This is designed to counteract premature termination of the localization routine due to e.g. flickering of the fluorophore target. When a number of integration roundtrips have passed without collecting the necessary number of photons (see Photon Limit) that in time equal the *maxOffTime* threshold, a break condition is triggered terminating the current attempt.

**Tip:** Exceeding the *maxOffTime* counts against the Number of Permitted Localization Attempts. It is encouraged to consider the *maxOffTime* as a sort of buffer to counteract unexpected events during tracking and increase consistency and track length as the SPT routine will not be so easily terminated. Keep in mind though that this may require additional post processing to remove e.g. mid-trace-particle-change events or longer lingering times after a particle has been lost and a new one found.

**Expected Input:** Floating-Point Number. The default “unspecified” equals $3ms$.
