## Supplementary Table 1 for "Parameter Optimization for Iterative MINFLUX Microscopy enabled Single Particle Tracking"

***Supplementary Table 1 – Complete Experimental Results Reveal Optimization Tradeoff Between Trackable Diffusion Rates and Localization Error for iMFX-enabled Single Particle Tracking*** – In addition to the ideal CRLB $\sigma_{CRLB}(\bar{r} = \bar{0})$ (see Supplementary Material of [7], p.11) we show the mean of the apparent lateral diffusion coefficients $D_{eMSD}$ (ensemble average) and $D_{tMSD}$ (time average), as well as the lateral localization error $\sigma_{eMSD}$ extracted using Optimal Least Squares Fitting (OLSF) on the ensemble average Mean-Squared-Displacement (*e*MSD) and time average Mean-Squared-Displacement (*t*MSD) curve respectively assuming momentary Brownian motion. Here $\eta$ refers to the number of cycles, $t_{loc}$ is the time-to-localization, $N$ is the number of photons, $N_{PL}$ is the *PL* and $\langle \dots \rangle$ the scale-appropriate average value. Shading indicates sample reference Dye on GUV-patch SLB (orange), QDs on GUV-patch SLB (green), QDs on Lipid-Deposition-SLB (blue) and the TIRF reference (lilac).

| PL | DMP | DT | $D_{eMSD}$ $[\mu m^2/s]$ | $D_{tMSD}$ $[\mu m^2/s]$ | $\frac{D_{eMSD}}{D_{tMSD}}$ | $\sigma_{eMSD}$ $[nm]$ | $\sigma_{CRLB}$ $[nm]$ | $\langle \eta \rangle$ | $\langle t_{loc} \rangle$ $[\mu s]$ | $\frac{\langle N \rangle}{\langle \eta \rangle N_{PL}}$ |
|---|---|---|---|---|---|---|---|---|---|---|
| 10 | 0 | 100 | 2.46 | 2.51 | 0.98 | 30.30 | 20.03 | 1 | 150 | 2.13 |
| 10 | **5** | 100 | 0.07 | 0.08 | 0.85 | 16.52 | 20.03 | 1 | 150 | 4.10 |
| 10 | **2** | 100 | 0.49 | 0.51 | 0.96 | 15.53 | 20.03 | 1 | 150 | 4.49 |
| 10 | **0** | 100 | 0.63 | 0.77 | 0.81 | 28.02 | 20.03 | 1 | 150 | 3.36 |
| **10** | 0 | 100 | 0.63 | 0.78 | 0.81 | 27.96 | 20.03 | 1 | 150 | 3.36 |
| **20** | 0 | 100 | 0.52 | 0.61 | 0.84 | 22.78 | 14.35 | 1 | 151 | 2.27 |
| **50** | 0 | 100 | 0.48 | 0.60 | 0.80 | 14.05 | 9.08 | 2 | 282 | 0.71 |
| **10** | 0 | 100 | 0.46 | 0.61 | 0.77 | 30.37 | 20.03 | 1 | 150 | 1.53 |
| **20** | 0 | 100 | 0.36 | 0.46 | 0.78 | 23.51 | 14.35 | 2 | 281 | 0.67 |
| **50** | 0 | 100 | 0.23 | 0.47 | 0.50 | 13.46 | 9.08 | 5 | 676 | 0.22 |
| 50 | 0 | **50** | 0.28 | 0.40 | 0.70 | 4.67 | 9.08 | 5 | 437 | 0.23 |
| 50 | 0 | **100** | 0.28 | 0.33 | 0.85 | 5.50 | 9.08 | 3 | 414 | 0.42 |
| 50 | 0 | **200** | 0.21 | 0.22 | 0.93 | 10.91 | 9.08 | 2 | 483 | 0.82 |
| **TIRF SPT** | | | 0.64 | 0.63 | 1.01 | 39.86 | – | 1 | 14925 | – |
